# Long-read DNA metabarcoding of ribosomal rRNA in the analysis of fungi from aquatic environments

**DOI:** 10.1101/283127

**Authors:** Felix Heeger, Elizabeth C. Bourne, Christiane Baschien, Andrey Yurkov, Boyke Bunk, Cathrin Spröer, Jörg Overmann, Camila J. Mazzoni, Michael T. Monaghan

## Abstract

DNA metabarcoding is now widely used to study prokaryotic and eukaryotic microbial diversity. Technological constraints have limited most studies to marker lengths of *ca.* 300-600 bp. Longer sequencing reads of several 5 thousand bp are now possible with third-generation sequencing. The increased marker lengths provide greater taxonomic resolution and enable the use of phylogenetic methods of classification, but longer reads may be subject to higher rates of sequencing error and chimera formation. In addition, most well-established bioinformatics tools for DNA metabarcoding were originally 10 designed for short reads and are therefore not suitable. Here we used Pacific Biosciences circular consensus sequencing (CCS) to DNA-metabarcode environmental samples using a *ca.* 4,500 bp marker that included most of the eukaryote ribosomal SSU and LSU rRNA genes and the ITS spacer region. We developed a long-read analysis pipeline that reduced error rates to levels 15 comparable to short-read platforms. Validation using fungal isolates and a mock community indicated that our pipeline detected 98% of chimeras *de novo* i.e., even in the absence of reference sequences. We recovered 947 OTUs from water and sediment samples in a natural lake, 848 of which could be classified to phylum, 486 to family, 397 to genus and 330 to species. By 20 allowing for the simultaneous use of three global databases (Unite, SILVA, RDP LSU), long-read DNA metabarcoding provided better taxonomic resolution than any single marker. We foresee the use of long reads enabling the cross-validation of reference sequences and the synthesis of ribosomal rRNA gene databases. The universal nature of the rRNA operon and our recovery of >100 25 non-fungal OTUs indicate that long-read DNA metabarcoding holds promise for the study of eukaryotic diversity more broadly.

## INTRODUCTION

DNA-metabarcoding is widely used in the study of microbial communities from all three major domains of life (Wurzbacher 2017), whereby one or more 30 marker regions in the genome are PCR-amplified and sequenced using a next-generation sequencing (NGS) platform. Reads are quality-filtered and sequences are clustered according to sequence similarity into putative taxa (Operational Taxonomic Units = OTUs). OTUs are then classified using marker-specific, and sometimes taxon-specific databases. DNA metabarcoding has 35 become a commonly used tool because it provides an estimate of biodiversity, including that of taxa that cannot be cultured, and identification relies on relatively stable genetic information rather than often variable and subtle phenotypic characters. Limitations of the method include the fact that marker regions and PCR primers must be selected *a priori* to detect the taxa of 40 interest, and that the variability of the marker region, and how well the taxa are represented within a given reference database, determine how well the members of an assemblage can be identified (Nilsson 2018).

There is a fundamental trade-of between using a marker that is conserved 45 enough to be amplified across a broad range of taxa, but variable enough to distinguish among closely related species. Marker length also has consequences for how many OTUs can be identified, and to what taxonomic resolution (Porras-Alfaro 2014). Shorter markers within a given locus may include less genetic variation than longer markers, reducing the ability to 50 distinguish closely related species (Singer 2016). One consequence is that highly variable regions are often used as DNA metabarcoding markers. While variable regions may increase taxonomic resolution in groups for which reference sequences are available, sequence homology can be difcult or impossible to establish. This precludes phylogeny-based analyses and can 55 result in the complete failure of classifying OTUs at any taxonomic level (Lindahl 2013).

More recent (i.e. third-generation sequencing) technologies can provide much longer (several kbp) sequencing reads (Goodwin 2016); however, their use in 60 studies of environmental samples remains limited. The few existing studies, using full-length (∼1.5 kbp) bacterial 16S (Franzén 2015, Schloss 2016, Singer 2016) and parts of the eukaryotic rRNA operon including ITS (up to 2.6 kbp) (Tedersoo 2017, Schlaeppi 2016), have reported increased taxonomic resolution. The Pacific Biosciences (PacBio) RSII platform generates reads of 65 >50 kbp by Single Molecule Real Time (SMRT) Sequencing. Single pass error rates of 13-15% (Goodwin 2016) limit their value in DNA metabarcoding because species identification is unreliable at those levels of uncertainty. However, the circular consensus sequencing (CSS) version of SMRT sequencing greatly reduces the error rate. In CSS, double stranded DNA amplicon 70 molecules are circularized by the ligation of hairpin adapters. The sequencing polymerase is then able to pass around the molecule and read the same insert multiple times (Travers 2010). The repeated reads of the same amplicon molecule, together with the random nature of sequencing error, can then be used to reduce the final error rate to <1% (Goodwin 2016) by generating 75 consensus sequences.

Beside the higher per base cost a primary reason why long-read approaches have not been applied to DNA metabarcoding is the fact that most of the existing bioinformatic tools have been optimized for the analysis of data from 80 short-read technologies (e.g., Illumina). It is thus unclear how well they will perform on PacBio CCS reads. Longer sequences have more errors because even high-quality reads with low error rates will accumulate more total errors as a function of length. The types of errors in PacBio reads also differ from that of short-read technologies, with CCS reads tending to have more insertions and deletions, compared to substitutions more common in short-read data. Schloss et al. (2016) explored the error profile and steps that can be taken when targeting the 16S for a bacterial mock community, and environmental samples. They found that the error rate of CSS reads of their longest amplicon (V1-V9) was only 0.68% and could be further reduced to 0.027% by pre-clustering at 99% similarity. Chimera formation rate may also be increased in longer markers since longer amplicons may suffer premature elongation terminations, leading to more possibilities for the resulting incomplete amplicons to act as primers in the next PCR cycle and thus more chimeras to be formed (see also Laver at al. 2016). Existing algorithms commonly used to detect chimeras are not optimized for long reads and may therefore fail to detect chimeras.

Fungi are ecologically important eukaryotes, having diverse roles in carbon and nutrient cycling, occupying a range of niches, including as decomposers, parasites and endophytes, and are ubiquitous in terrestrial and aquatic habitats alike (e.g. Tedersoo 2014, Wurzbacher 2016). Microbial fungal communities are increasingly studied with DNA metabarcoding (e.g., Roy 2017), taking advantage of the increased detection of taxa without the need to culture and the reduced cost of sequencing that has permitted ever deeper read depth The broad phylogenetic diversity of fungi has the consequence that fungal DNA metabarcoding studies typically use markers that vary depending on the taxonomic group of interest and the resolution desired. Different regions of the eukaryotic rRNA operon have been widely utilized for barcoding fungi due to its universality, and the fact that short stretches have been able to provide reasonable power for fungal identification. Within this region, the most commonly applied barcode is the internal transcribed spacer (ITS) (Schoch 2012). This comprises the ITS1, the 5.8S rRNA gene and the ITS2, and depending on the lineage, varies from 300 to 1,200 bp in length. In fungal DNA metabarcoding, the ITS2 region is widely used to assess fungal diversity in environmental samples (Blaalid 2013, Kõljalg 2013); however, it is not as successful in identifying taxa as the full length ITS (Tederso 2017). For early diverging fungal lineages, such as those found in many aquatic habitats (Monchy 2011, Wurzbacher 2016, Rojas-Jimenez 2017), sequences from the small subunit (SSU) rRNA gene (18S) can provide afliation of higher taxonomic ranks, but are often not variable enough to distinguish among species (Cole 2014). The LSU region has higher variability, and therefore resolution, than the SSU, and is often used for identification of specific fungal groups (e.g. Glomeromycota and Chytridiomycota) lacking ITS reference sequences. Databases have been established for all three different markers, e.g. UNITE for ITS (Kõljalg 2013), SILVA for SSU (Quast 2013), and RDP for LSU (Cole 2014). Nevertheless, database coverage remains poor for several fungal lineages, for example Glomeromycota (Ohsowski 2014), Chytridridiomycota (Frenken 2017), and Cryptomycota, and for species from less-studied habitats such as aquatic, indoor, and marine environments.

We examined fungal diversity of field-collected samples from a temperate lake using SMRT CCS of a long (*ca.* 4,500 bp) DNA metabarcode that included the three major regions of the eukaryotic rRNA operon (SSU, ITS, LSU) in a single sequencing read. We first sequenced cultured isolates comprising a broad phylogenetic range and a mock community to derive rates of sequencing error and chimera formation. We then developed a new bioinformatics pipeline designed for full length rRNA operon amplicons. We found error rates to be comparable to short-read approaches after filtering with our pipeline, and chimera-formation rates to be comparable to those found in studies with shorter amplicons. We identified 947 OTUs from environmental samples, 848 of which could be classified to phylum, 486 to family, 397 to genus and 330 to species. By allowing for the simultaneous use of three databases, long-read DNA metabarcoding provided much better taxonomic resolution than possible with a single-marker, single-database approach. The universal nature of the rRNA operon and our recovery of >100 non-fungal OTUs indicate that long-read DNA metabarcoding holds promise for future studies of eukaryotic diversity in general.

## METHODS

### Isolates, Mock community, and Environmental samples

Isolates of sixteen fungal species (Table 1) were combined to form a mock community. This community was used to test PCR and library preparation protocols that were later applied to environmental samples, and to quantify the efciency of *de novo* and reference-based chimera detection in our long-read bioinformatics pipeline described below. Environmental samples were collected from Lake Stechlin, an oligo-mesotrophic lake in North-East Germany (53.143° N 13.027° E) in October 2014. Littoral water samples (30 L total) were collected and pooled from surface water in the shallow zone along three 10 m transects, located within 5 m of the lake shore or reed belt. Pelagic water samples (30 L total) were collected from the deeper zone of the lake by pooling samples taken at multiple depths (0-65 m) at one point, using a Niskin-bottle (Hydro-Bios, Kiel, Germany). A subsample (2 L) of each (littoral and pelagic) was filtered through 0.22-µmm Sterivex filters (Merck Millipore, Darmstadt, Germany) using a peristaltic pump (GT-EL2 Easy Load II, UGT, Müncheberg, Germany). Excess water was expelled using a sterile syringe and parafilm used to seal the ends. Sediment samples were collected from four locations in each zone (littoral, pelagic) using a PVC sediment corer (63 mm diameter) on a telescopic bar (Uwitec, Mondsee, Austria). The uppermost 2 cm from each sediment core were pooled in the field and divided into 2 ml subsamples for storage. Sterivex filters and sediment subsamples were frozen in liquid Nitrogen in the field and returned to the laboratory for long-term storage at −80°C.

**Table 1.**
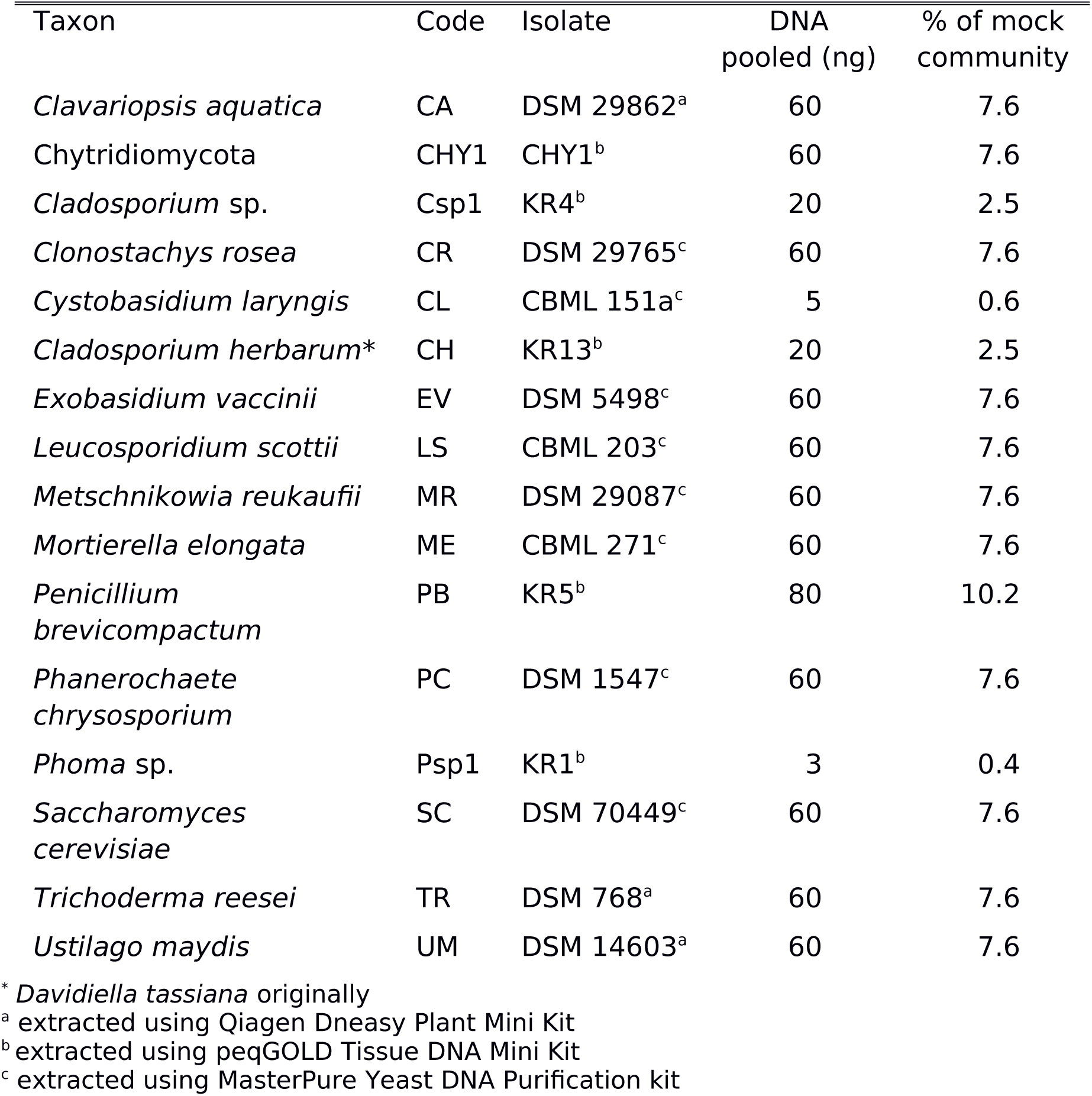
Isolates used and their contribution to the mock community.

| Taxon | Code | Isolate | DNA<br>pooled (ng) | % of mock<br>community |
| --- | --- | --- | --- | --- |
| *Clavariopsis aquatica* | CA | DSM 29862<sup>a</sup> | 60 | 7.6 |
| Chytridiomycota | CHY1 | CHY1<sup>b</sup> | 60 | 7.6 |
| *Cladosporium* sp. | Csp1 | KR4<sup>b</sup> | 20 | 2.5 |
| *Clonostachys rosea* | CR | DSM 29765<sup>c</sup> | 60 | 7.6 |
| *Cystobasidium laryngis* | CL | CBML 151a<sup>c</sup> | 5 | 0.6 |
| *Cladosporium herbarum* * | CH | KR13<sup>b</sup> | 20 | 2.5 |
| *Exobasidium vaccinii* | EV | DSM 5498<sup>c</sup> | 60 | 7.6 |
| *Leucosporidium scottii* | LS | CBML 203<sup>c</sup> | 60 | 7.6 |
| *Metschnikowia reukaufii* | MR | DSM 29087<sup>c</sup> | 60 | 7.6 |
| *Mortierella elongata* | ME | CBML 271<sup>c</sup> | 60 | 7.6 |
| *Penicillium*<br>*brevicompactum* | PB | KR5<sup>b</sup> | 80 | 10.2 |
| *Phanerochaete*<br>*chrysosporium* | PC | DSM 1547<sup>c</sup> | 60 | 7.6 |
| *Phoma* sp. | Psp1 | KR1<sup>b</sup> | 3 | 0.4 |
| *Saccharomyces*<br>*cerevisiae* | SC | DSM 70449<sup>c</sup> | 60 | 7.6 |
| *Trichoderma reesei* | TR | DSM 768<sup>a</sup> | 60 | 7.6 |
| *Ustilago maydis* | UM | DSM 14603<sup>a</sup> | 60 | 7.6 |

\* *Davidiella tassiana* originally
<sup>a</sup> extracted using Qiagen Dneasy Plant Mini Kit
<sup>b</sup> extracted using peqGOLD Tissue DNA Mini Kit
<sup>c</sup> extracted using MasterPure Yeast DNA Purification kit

### DNA extraction

Genomic DNA was extracted from fungal isolates using three different methods (Table 1, see also Supp. Info 1). Environmental DNA was extracted from water and sediment samples using a modified phenol-chloroform method (after Nercessian 2005). Frozen Sterivex cartridges were broken open and sterilized forceps were used to transfer half of the fragmented filter into each of two 2-ml tubes. Sediment samples were thawed and aliquoted into two 2-ml tubes, each containing 200 mg. Beads (0.1 and 1.0 mm zirconium, and 3x 2.5mm glass beads, Biospec, Bartlesville, USA) were added to 0.3 volume of the tube. For cell lysis and extraction, the following reagents were added: 0.6 ml CTAB extraction buffer (5% CTAB-120 mM phosphate buffer), 60 µml 10% sodium dodecyl sulfate, 60 µml 10% N-lauroyl sarcosine, followed by 0.6 ml of phenol:chloroform-isoamyl alcohol (25:24:1). Samples were vortexed immediately to homogenise and then ground for 1.5 min at 30 Hz (Retsch mill, Retsch GmbH, Haan, Germany) with short breaks for cooling on ice. Samples were incubated for 1 hr at 65 °C, with occasional mixing, and then centrifuged at 17,000 g for 10 minutes. The upper aqueous phase was transferred to a new tube and mixed with an equal volume of chloroform-isoamyl alcohol (24:1), centrifuged at 17,000 g for 10 min and the upper aqueous phase transferred to a new tube. Nucleic acids were precipitated with 2 volumes of PEG/NaCl (30% PEG 6000 in 1.6 M NaCl) for 2 h. Samples were centrifuged at 16,000 g for 45 min, and the supernatant discarded. The nucleic acid pellet was washed twice by the addition of 1 ml ice-cold 70% ethanol, centrifuged at 17,000 g for 15 min, and the supernatant discarded and following removal of ethanol traces, eluted in 50 µml nuclease-free water. Subsamples were pooled to give 100 µml nucleic extract per sample. RNA was removed by the addition of 0.5 µml (5 µmg) RNase A (10 mg/ml DNase and protease free, ThermoFisher Scientific, Waltham, US) to 80 µml of the pooled sample, incubated at 37 °C for 30 min, and cleaned using the PowerClean Pro DNA Clean-Up kit (MoBio Laboratories, Carlsbad, USA). DNA was quantified in triplicate using a Qubit HS dsDNA Assay (Invitrogen, Carlsbad, USA) and gel-checked for quality.

### PCR and chimera formation tests

Approximately 4,500 bp of the eukaryotic rRNA operon (Fig 1), including SSU, ITS1, 5.8S, ITS2, and LSU (partial) regions, was PCR-amplified using the primers NS1_short and RCA95m (C. Wurzbacher, unpublished). NS1_short (5’-CAGTAGTCATATGCTTGTC-3’) was modified from White et al. (1990) by shortening to remove several major mismatches to fungal groups. RCA95m (5’-ACCTATGTTTTAATTAGACAGTCAG-3’) was modified from R78 (Wurzbacher 2014). Symmetric (reverse complement) 16-mer barcodes (Supplemental Table 1) were added to the 5’ ends of primers following the PacBio manufacturer’s guidelines on multiplexing SMRT sequencing.

**Figure 1:**
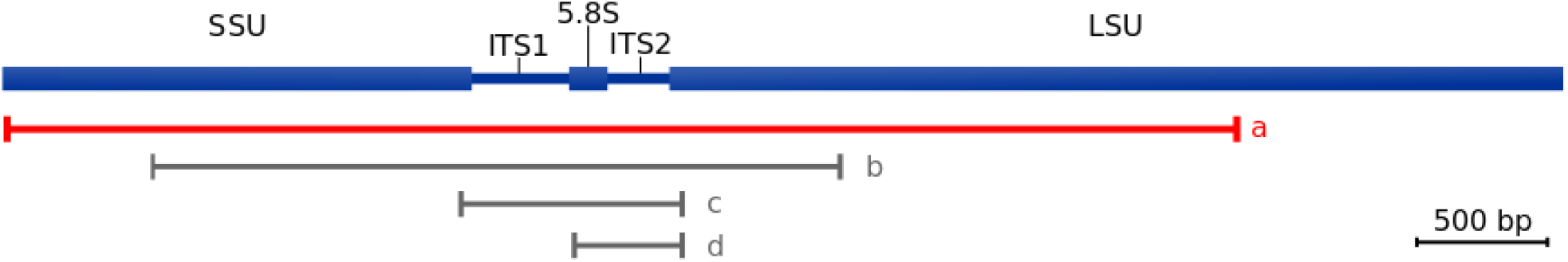
Region of the eukaryotic rRNA operon covered by the primer pair used in this studied (a) compared to the primer pair SSU515Fngs-TW13 used by Tedersoo et al. 2017 (b), the widely used (e.g. Schoch 2012) primer pairs ITS5-ITS4 (c) and ITS3-ITS4 (d)

We aimed to minimize chimera formation by minimizing the number of PCR cycles performed per sample. Cycle numbers were chosen after amplifying all samples with a variable number of cycles (13-30) and identifying the exponential phase of PCR (Lindahl 2013) according to band visibility on an agarose gel. Based on these results, we used 15-20 cycles to amplify isolates (3-8 ng template DNA), 13-30 cycles for mock community samples (2-20 ng), and 22-26 cycles for environmental samples (10 ng). Barcodes were allocated to the different PCR conditions tested as shown in supplemental Tables 2 and 3. All standard PCRs were conducted in 25 µml reactions using 0.5 µml Herculase II Fusion enzyme (Agilent Technologies, Cedar Creek, USA), 5 µml of 5x PCR buffer, 0.62 µml each primer (10 uM), 0.25 µml dNTPs (250 mM each), 0.3 µml BSA (20mg/ml BSA, ThermoFisher Scientific, Waltham, US) on a SensoQuest labcycler (SensoQuest Gmbh, Göttingen, Germany) with 2 min denaturation at 95 °C, 13-30 cycles (see above) of 94 °C for 30 sec, 55 °C for 30 sec and 70 °C for 4 min, and a final elongation at 70 °C for 10 min. Multiple PCR reactions (up to 50) were required for each environmental sample to ensure sufcient product for library preparation (1 µmg purified PCR product). We also included a two-step emulsion PCR (emPCR) of the mock community in order to test whether emPCR could reduce chimera formation rate by the physical isolation of DNA template molecules (Boers 2015). The Micellula DNA Emulsion kit (Roboklon GmbH, Berlin) was used for a two-step PCR: a first amplification of 25 cycles, with 2µml of the cleaned template used in a second 25 cycle PCR. For further details see supplemental info 2.

### Library preparation and Sequencing

Replicate PCRs were pooled back to sample level, and products were cleaned with 0.45 x CleanPCR SPRI beads (CleanNa, Waddinxveen, Netherlands), pre-cleaned according to PacBio specifications (C. Koenig, pers. comm.), quantified twice using a Qubit HS dsDNA Assay, and quality-checked on an Agilent® 2100 Bioanalyzer System (Agilent Technologies, Santa Clara, USA). Samples were then pooled into libraries (as described in supplemental Table 3) before being quality-checked on an Agilent® 2100 Bioanalyzer following PacBio guidelines (Pacific Biosciences, Inc., Menlo Park, CA, USA) for amplicon template library preparation and sequencing.

SMRTbell™ template libraries were prepared according to the manufacturer’s instructions following the Procedure & Checklist – Amplicon Template Preparation and Sequencing (Pacific Biosciences). Brieey, amplicons were end-repaired and ligated overnight to hairpin adapters applying components from the DNA/Polymerase Binding Kit P6 (Pacific Biosciences). We included enough DNA from each sample to obtain the required library concentration (37 ng µml^-1^) for end-repair. Reactions were carried out according to the manufactureŕs instructions. Conditions for annealing of sequencing primers and binding of polymerase to purified SMRTbell™ template were assessed with the Calculator in RS Remote (Pacific Biosciences). SMRT sequencing was carried out on the PacBio *RSII* (Pacific Biosciences) taking one 240-minutes movie.

In total, we ran 8 libraries and 27 SMRT cells. Three of the isolates (*Trichoderma reesei, Clonostachys rosea,* and a species belonging to the phylum Chytridiomycota) were sequenced on one SMRT cell to test the protocol for CCS. The remaining 13 isolates and one of the mock community conditions (30 PCR cycles) were prepared as part of the libraries containing the environmental samples (Supplemental Table 3), which were each run on three SMRT cells. Mock community samples and the emPCR sample were pooled in equimolar ratio and sequenced using two SMRT cells.

Demultiplexing and extraction of subreads from SMRT cell data was performed applying the RS_ReadsOfInsert.1 protocol included in SMRTPortal 2.3.0 with minimum 2 full passes and minimum predicted accuracy of 90%. Barcodes were provided as FASTA files and barcode extraction was performed in a symmetric manner with a minimum barcode score of 23 within the same protocol. Mean amplicon lengths of 3800 – 4500 kbp were confirmed. Demultiplexed reads were downloaded from the SMRT Portal as fastq files for further analysis.

### Long-read metabarcoding pipeline

We developed an analysis pipeline for PacBio CCS reads using the python workeow engine snakemake (version 3.5.5, Köster & Rahmann 2012). Our pipeline combines steps directly implemented in python with steps that use external tools. The implementation is available on github (https://github.com/f-heeger/long_read_metabarcoding) and parameters used for the external tools can be found in the supplemental methods (supplemental info 3).

#### Read Processing stage

Reads longer than 6,500 bp were excluded to remove chimeric reads formed during adapter ligation and reads containing double-inserts due to failed adapter recognition during the CCS generation. Reads shorter than 3,000 bp were removed to exclude incompletely amplified sequences and other artifacts. Reads were then filtered by a maximum mean predicted error rate of 0.004 that was computed from the Phred scores. Reads with local areas of low quality were removed if predicted mean error rate was > 0.1 in any sliding window of 8 bp. cutadapt (version 1.9.1, Martin 2011) was used to remove forward and reverse amplification primers and discard sequences in which primers could not be detected. Random errors were reduced by pre-clustering reads from each sample at 99% similarity using the cluster_smallmem command in vsearch (version 2.4.3, Rognes 2016). Reads were sorted by decreasing mean quality prior to clustering to ensure that high quality reads were used as cluster seeds. vsearch was configured to produce a consensus sequence for each cluster.

#### OTU clustering and classification stage

Chimeras were detected and removed with the uchime_denovo command in vsearch. Based on tests using mock community samples (see below), we determined this was a suitable method of chimera detection following the read processing stage (above). Only sequences that were classified as non-chimeric were used for further analysis. The rRNA genes (SSU, LSU, 5.8S) and internal transcribed spacers (ITS1, ITS2) in each read were detected using ITSx (version 1.0.11, Bengtsson-Palme 2013). To generate OTUs, the ITS region (ITS1, 5.8S, ITS2) was clustered using vsearch at 97% similarity. SSU and LSU sequences were then placed into clusters according to how their corresponding ITS was clustered. OTUs were taxonomically classified using the most complete available database for each marker. For the ITS we used the general FASTA release of the UNITE database (version 7.1, 20.11.2016, only including singletons set as RefS, Kõljalg 2013); for the SSU we used the truncated SSU release of the SILVA database (version 128, Quast 2013), excluding database sequences with quality scores below 85 or Pintail chimera quality below 50; and for the LSU we used the RDP LSU data set (version 11.5, Cole 2014). The ITS, SSU and LSU regions of the representative sequence of each OTU were locally aligned to the database using lambda (version 1.9.2, Hauswedell 2014). For LSU and SSU the alignment parameters had to be modified to allow for longer alignments (see supplemental info 3). From the alignment results, a classification was determined by filtering the best matches and generating a lowest common ancestor (LCA) from their classifications as follows. For each query sequence, matches were filtered by a maximum e-value (10^-6^), a minimum identity (80%) and a minimum coverage of the shorter of the query or database sequence (85%). For the SSU and LSU, non-overlapping matches between each query and database sequence were combined. For each query sequence, a cutoff for the bit score was established representing 95% of the value for the best match, above which all matches for that given sequence were considered. For the SSU and LSU, bit scores were normalized by the minimum length of query and database sequences to account for the varying lengths of database sequences. To determine the LCA from the remaining matches, their classifications were compared at all levels of the taxonomic hierarchy starting at kingdom (highest) and ending at species (lowest) level. For each OTU, the classifications of all matches at a given taxonomic rank were compared and if >90% of them were the same then this was accepted. If <90% were the same then the OTU remained unclassified at this and all lower ranks.

### Error rates based on isolate sequences

Isolate sequences were processed using the Read Processing stage of the pipeline (described above) in order to generate error-corrected consensus sequences from pre-clusters. The consensus sequences of the largest pre-cluster for each isolate were > 99 % identical to the Sanger sequencing data obtained from the same isolate (not shown), with most differences found in bases that were of low quality in the Sanger sequence data. We therefore used the consensus sequence of the largest cluster for each isolate as a reference for that species in all further analysis. CSS reads from each isolate were then aligned with the respective consensus sequence using blasr (github comit 16b158d, Chaisson & Tesler 2012) to estimate error rates of CCS reads. Sequences after filtering steps were also compared in order to estimate remaining errors.

### Evaluating chimera detection

*De novo* and reference-based chimera classifications were compared as a way of estimating the reliability of *de novo* chimera calls. The CCS reads from the mock community samples were tested for chimeras with vsearch once in *de novo* mode (uchime_denovo) and once with a reference-based approach (uchime_ref). For the *de novo* approach, reads were processed with the Read Processing stage of the pipeline (above) to generate error-corrected sequences from pre-clusters. Cluster sizes resulting from the pre-clustering step were used as sequence abundances. For the reference-based approach, a reference file was created from the consensus sequence of the largest cluster for each isolate sample. A random subset of reads (100 sequences, 1.3% of the data) was generated from the mock community sample with the highest chimera rate and the most reads (30 PCR cycles). The subset of reads was aligned to the consensus sequences from the isolate samples and visually inspected for chimeras in Geneious (version 7.1.9, Kearse et al. 2012). These “manual” chimera calls were then used to verify reference-based chimera classifications for these reads. Chimeras identified by the reference-based approach were used to compute the chimera formation rate under different PCR conditions.

### Mock community classification

We tested classification with the DNA metabarcoding pipeline using the mock community sample with the most reads (30 PCR cycles). In the pipeline, chimeras were classified *de novo* and OTU classification was performed using the public databases. We manually classified the same OTUs using consensus sequences from our isolate samples as reference. For each read, chimeras were detected with a reference-based approach using vsearch and the classification of the read was determined by mapping reads to the isolate sample sequences with blasr. To better understand the resolution that can be expected from the different regions of the rRNA operon, each region (SSU, ITS1, 5.8S, ITS2, LSU) was clustered independently. Chimeras were first removed using the reference-based approach with our isolate sequences as references. The different regions in each read were separated with ITSx, dereplicated and clustered at 97%.

### Environmental community classification

Sequences from the environmental samples from Lake Stechlin were processed with the full rRNA metabarcoding pipeline described above. Chimeras were detected using the *de novo* approach, which we conclude provides a very good diagnosis of chimeras based on our validation using the mock community to compare *de novo* and reference-based approaches (see Results). The resulting classifications obtained with SSU, ITS, and LSU markers were then compared at each taxonomic level. OTUs with only one read (singletons) were excluded from this comparison.

## RESULTS

Sequencing resulted in a total number of 235,827 CCS reads, which were submitted to the NCBI Sequence Read Archive (SRR6825218 - SRR6825222). 218,032 of these reads were within the targeted size range of 3,000 – 6,500 bp (Table 2). After stringent filtering using average- and window-quality criteria, 70,308 reads remained that contained an identifiable amplification primer sequence (Table 2). Pre-clustering of isolate samples with the metabarcoding pipeline resulted in one large (> 80 reads) pre-cluster for each sample. Besides these big clusters, six samples had additional very small (< 3 reads) clusters. For isolates sequenced on two different SMRT-cells, consensus sequences of the large pre-clusters were identical across cells except for *Saccharomyces cerevisiae* where a T homopolymer in the ITS2 was 6 bases long in one consensus and 7 in the other. Consensus sequences of large clusters were used as reference for further analysis and submitted to gene bank (MH047187 - MH047202). The mean sequencing error rate of quality-filtered CCS reads, based on comparison to the consensus sequences of the large clusters (taken to be our reference for each isolate), was 0.223% (SD 1.558%). Deletions were by far the most common error (0.179%), with insertions and substitutions much lower (Table 3).

**Table 2.** Number of sequencing reads remaining after each step in the bioinformatics pipeline for each sample type.

| Analysis step | Isolates | Mock community | Environmental samples | Total |
| --- | --- | --- | --- | --- |
| Raw CCS | 50,118 | 59,683 | 126,026 | 235,827 |
| Length-filtered | 47,766 | 52,871 | 117,395 | 218,032 |
| Average quality-filtered | 20,009 | 15,686 | 48,778 | 84,473 |
| Window quality-filtered | 17,675 | 10,927 | 43,385 | 71,987 |
| Primer-filtered | 17,380 | 10,559 | 42,369 | 70,308 |

**Table 3.** Error rates in CSS reads computed by mapping to consensus sequences of isolates.

| Analysis step | Substitutions mean (SD) | Insertions mean (SD) | Deletions mean (SD) | Total mean (SD) |
| --- | --- | --- | --- | --- |
| Raw CSS | 0.0450%<br>(0.1291%) | 0.3102%<br>(0.6076%) | 0.8671%<br>(1.1980%) | 1.2224%<br>(1.5577%) |
| Filtered | 0.0076%<br>(0.0264%) | 0.0364%<br>(0.0466%) | 0.1790%<br>(0.1575%) | 0.2230%<br>(0.1639%) |

### Chimera formation and detection

Using reference-based chimera detection in the mock community, chimera formation rate (i.e. sequences classified as chimeras or as unsure) rose from 3% of sequences at 13-18 PCR cycles to 16% at 30 cycles (Fig. 2). The emPCR (25 cycles) resulted in 4% of sequences classified as chimeric (Fig. 2), compared to 14% for 25 cycles under standard PCR conditions. Template DNA amounts played no measurable role in chimera formation rate, with 2, 8 and 20 ng of DNA all resulting in <2% chimeric sequences (18 cycles). Manual inspection of 100 randomly chosen isolate sequences classified 16 of these as chimeras. Reference-based detection identified 15 of these as chimeric and one as “suspicious”. Of the 84 confirmed as non-chimeric by manual inspection, the reference-based algorithm classified 82 (97.6%) as non-chimeric and 2 as “suspicious”. *De novo* chimera detection (i.e., in the absence of a reference) classified 98.5% of the reads in the sample in the same way as using the reference-based approach.

**Figure 2:**
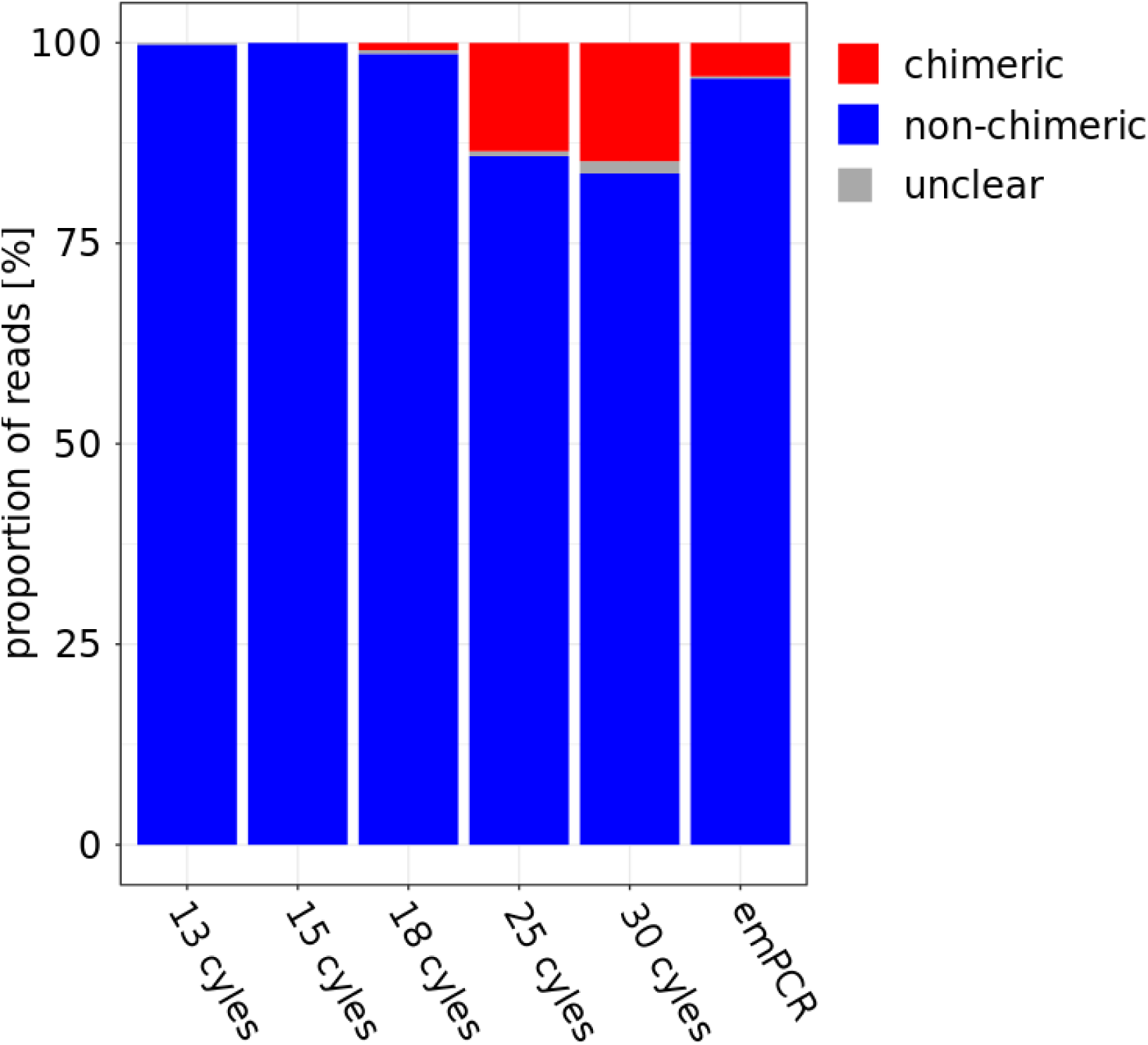
Chimera calls by vsearch with reference-based approach for different PCR conditions. Reads are classified as “chimeric” (red), “non-chimeric“ (blue) or in edge cases as “unclear” (gray).

### Mock community classification

The fve marker regions (SSU, ITS1, 5.8S, ITS2, LSU) clearly distinguished 8 of the 14 isolates we could recover within the mock community, but revealed cases of intra-specific variation as well as overlap among recognized species (Fig. 3). Seven species were clearly distinguished at all five markers, i.e. formed a single cluster for each region (Fig. 3). *Metschnikowia reukaufii* produced multiple clusters for ITS1 and ITS2, as expected based on previous reports of extraordinarily high rRNA operon variation in this genus (Sipiczki 2013, Lachance 2003). *Clavariopsis aquatica* and *Phoma* sp. were separated by all regions except SSU. *Trichoderma reesei* and *Clonostachys rosea* were separated by ITS1, ITS2, and LSU but not with SSU and 5.8S genes. *Cladosporium herbarum* and *Cladosporium* sp. were differentiated only with the ITS2, although one of the two clusters was mixed (Fig. 3). OTU clustering resulted in 16 non-singleton OTUs. Twelve OTUs consisted of sequences from one species as well as a few chimeric sequences, one contained sequences from *Cladosporium herbarum* and the other *Cladosporium* sp., and three smaller OTUs were entirely made up of chimeric sequences (Table 4). *Mortierella elongata* and *Cystobasidium laryngis* did not appear in any OTUs, although we did observe low read abundance (<10 reads) of these species prior to quality filtering.

**Figure 3:**
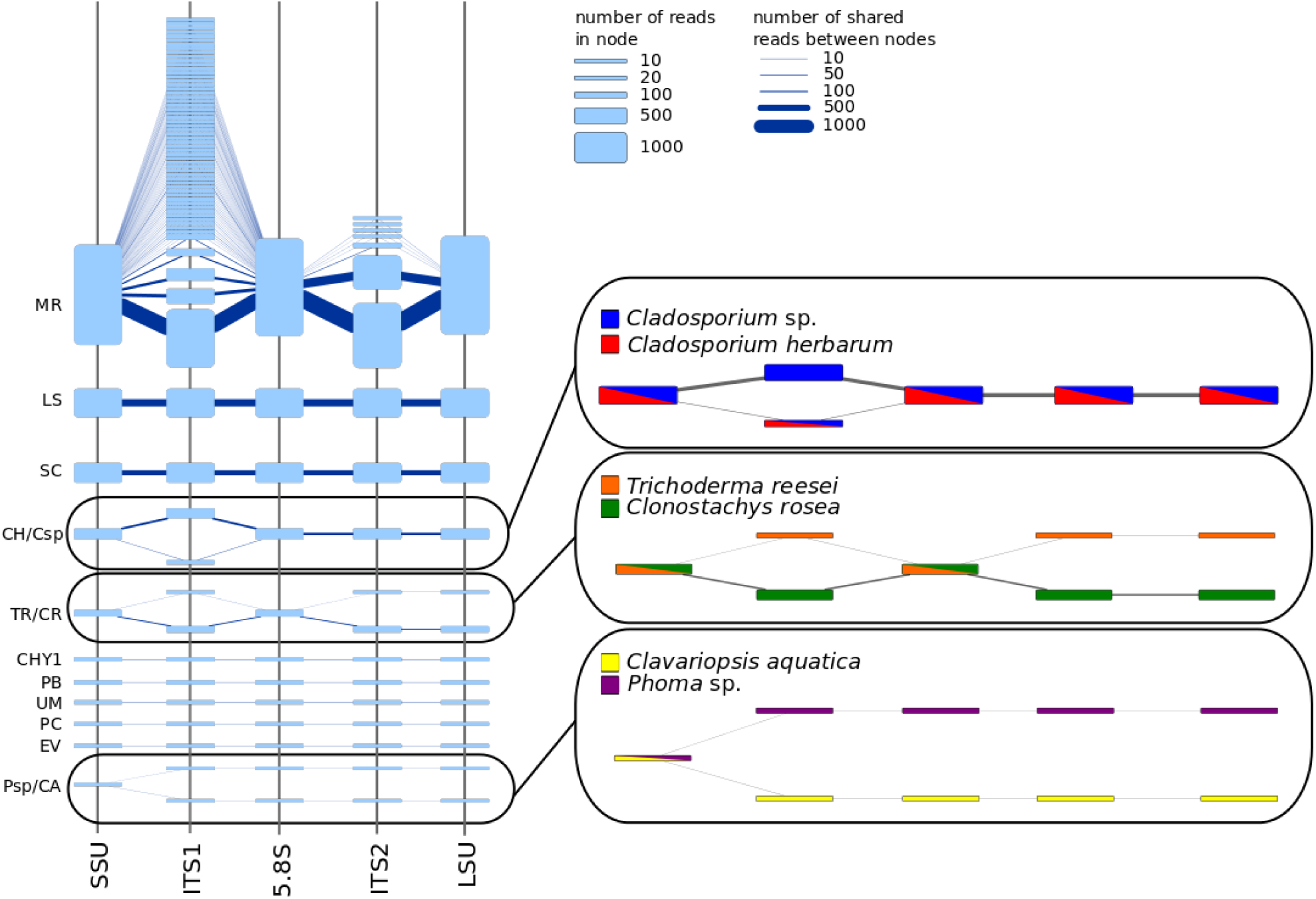
Resolution of different regions of the rRNA operon for our mock community. Each node represents a cluster and each edge between two clusters represents shared reads between the clusters. Node height and edge thickness is proportional to read number. Nodes and edges with less than 3 reads are not shown. Identification codes are given in Table 1. Components with multiple species are shown in detail on the right. Nodes are colored by species appearing in them. The graph was initially created with Cytoscape (version 3.2.1, Shannon 2003) and manually adapted for better readability.

**Table 4.** Mock-community OTU classification with our analytical pipeline. Manual classifications were made by comparison to full-length reference sequences. rRNA gene region classifications were made based on reference sequences in SILVA (SSU), UNITE (ITS) and RDP (LSU) databases. Size indicates the number of reads.

| OTU | Size | Classification method |  |  |  |
| --- | --- | --- | --- | --- | --- |
|  |  | Manual | SSU | ITS | LSU |
| 11 | 6 | *Clavariopsis aquatica* | Pleosporales (Order) | *Clavariopsis aquatica* | Pleosporales (Order) |
| 6 | 44 | Chytridiomycota | Chytridiomycetes (Class) | Globomyces (Genus) | Rhizophydium (Genus) |
| 4 | 344 | *Cladosporium* sp. +<br>*Cladosporium herbarum* | Cladosporium (Genus) | Cladosporium (Genus) | Davidiella (Genus) |
| 5 | 140 | *Clonostachys rosea* | Hypocreales (Order) | Bionectriaceae (Family) | Hypocreales (Order) |
| 1 | 4165 | *Metschnikowia reukaufii* | Saccharomycetales (Order) | *Metschnikowia cibodasensis* | *Metschnikowia bicuspidata* |
| 2 | 1096 | *Leucosporidium scottii* | Basidiomycota (Phylum) | Leucosporidiaceae (Family) | *Leucosporidium* (Genus) |
| 3 | 719 | *Saccharomyces cerevisiae* | Saccharomycetaceae<br>(Family) | *Saccharomyces* (Genus) | *Saccharomyces* (Genus) |
| 7 | 37 | *Penicillium brevicompactum* | Trichocomaceae (Family) | *Penicillium* (Genus) | Fungi (Kingdom) |
| 8 | 34 | *Ustilago maydis* | Ustilaginaceae (Family) | Ustilaginaceae (Family) | *Ustilago maydis* |
| 9 | 20 | *Exobasidium vaccinii* | Exobasidiales (Order) | *Exobasidium vaccinii* | Exobasidium (Genus) |
| 10 | 21 | *Phanerochaete chrysosporium* | Agaricomycetes (Class) | *Phanerochaete* sp. | Agaricomycetes (Class) |
| 12 | 5 | *Phoma* sp. | Pleosporales (Order) | Pleosporales Incertae sedis (Family) | Didymellaceae (Family) |
| 13 | 5 | *Trichoderma reesei* | Hypocreaceae (Family) | Trichoderma (Genus) | Hypocreaceae (Family) |
| 14 | 3 | chimeric | Saccharomycetales (Order) | Nectriaceae (Family) | *Metschnikowia bicuspidata* |
| 16 | 3 | chimeric | Saccharomycetales (Order) | *Metschnikowia cibodasensis* | unknown |
| 17 | 9 | chimeric | Saccharomycetales (Order) | *Metschnikowia cibodasensis* | Bionectria (Genus) |

OTUs were classified to varying taxonomic ranks by the three different genetic markers (Table 4). The SSU gene provided mainly order- and family-level classifications, the ITS region provided family-to species-level classifications, and the LSU gene provided genus-level classifications in some cases and higher level classifications in others. The *Metschnikowia reukaufii* OTU was classified to different species by ITS (*M. cibodasensis*) and LSU (*M. bicuspidata*). Different genus-level classifications by ITS and LSU for the Chytrid species were the result of different taxonomies used in the UNITE and the RDP databases. The best match in both databases was *Globomyces pollinis-pini*, but the higher classification at higher ranks differs among the databases. Similar discrepancies caused by differences in database taxonomy also occurred for some of the other species. Other than that classifications by all three markers were consistent with each other and with the manual classification.

### Environmental community classification

OTU clustering of the environmental samples produced 947 non-singleton OTUs (supplemental table 4), of which 799 (84%) were classified as fungi by at least one of the three markers (SSU, ITS, LSU). The SSU database also allowed identification of non-fungal sequences, and 112 OTUs were assigned to Metazoa, 10 to Discicristoidea, 2 to Stramenopiles, 2 to Alveolata and 1 to Chloroplastida. The 200 most abundant fungal OTUs (91% of fungal reads; 61% of total reads) were consistently classified to phylum level by all three markers except for 9 cases in which SSU and LSU gave different classifications for the same OTU (Fig. 4). There were no coneicts between SSU and ITS, although the SILVA and UNITE databases use different names for the phylum Cryptomycota/Rozellomycota (Fig. 4). Classification at the phylum level was most successful with SSU (188 reads, i.e., 94% of the 200 most abundant fungal OTUs). Fewer OTUs were classified to phylum with LSU (126, 63%) and ITS regions (36, 18%). Classification to the species level was most successful with LSU (55, 27.5%) and less successful for ITS (20, 10%) and SSU (13, 6.5%) (Fig. 4).

**Figure 4:**
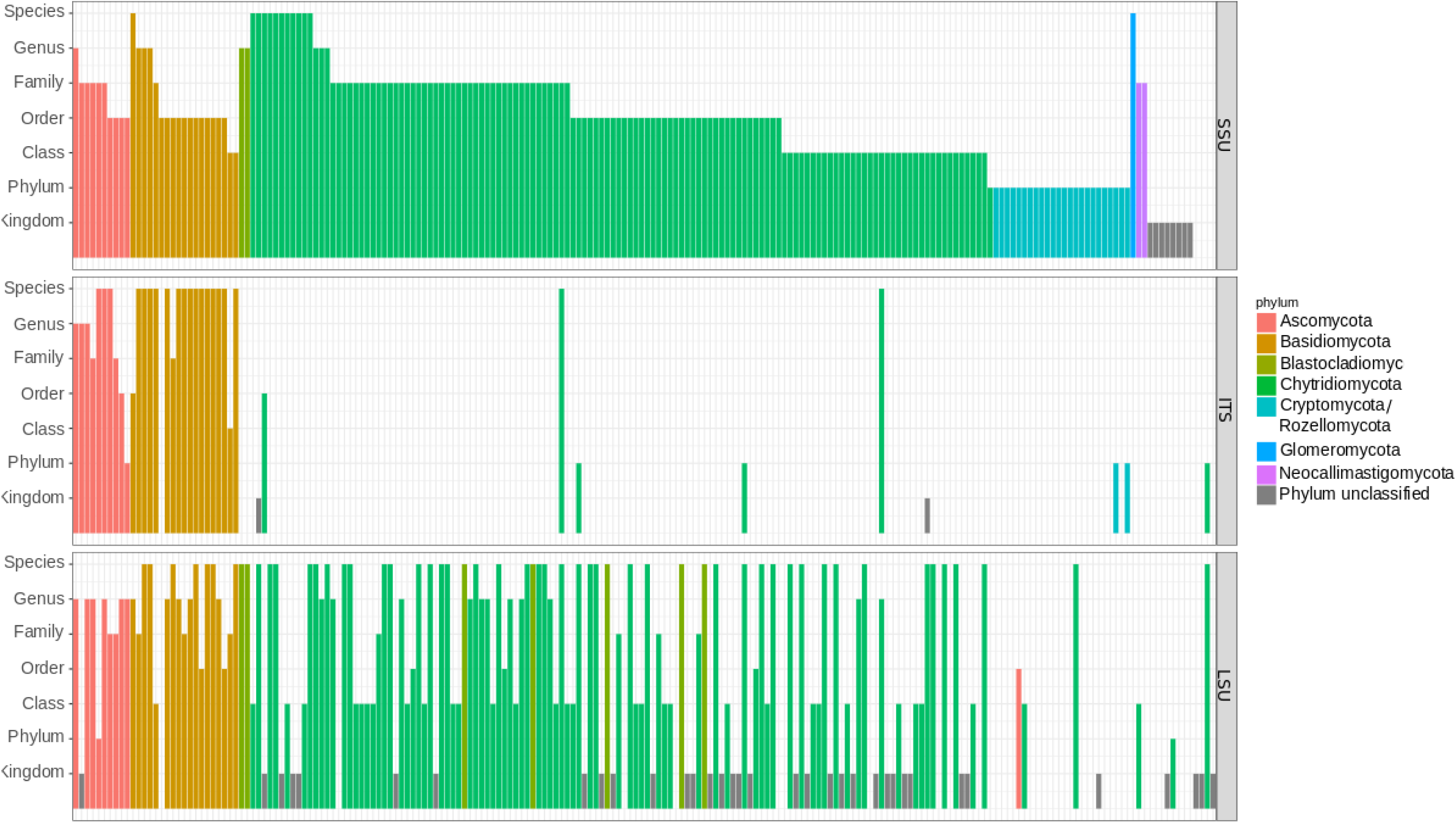
Classification specificity of the 200 most abundant fungal OTUs for the three different regions (SSU, ITS, LSU). The three rows give classifications by the three different regions. Each OTUs classification is given by a bar in each row. The height of the bar represents level of classification. Bars are colored by phylum.

Extended to all 947 OTUs, the results were similar. SSU provided the most classifications, especially for higher taxonomic ranks, and ca. 20% of these were classified the same using the ITS (Fig. 5A) and ca. 66% were classified the same by LSU (Fig. 5B). ITS classifications matched those of SSU (Fig. 5C) and LSU (Fig. 5D) at ranks from kingdom to class. At family, genus and species rank, most OTUs that were classified by ITS were not classified by SSU (Fig. 5C) and many were classified differently by LSU (Fig. 5D). At higher taxonomic rank (kingdom to class), OTUs classified by LSU were classified the same way as by SSU. But more than 50% were either not assigned to any taxon or were classified differently by SSU at lower ranks (order to species; Fig. 5E). Most OTUs classified by the LSU were not classified by ITS at kingdom to class ranks (> 60%), although those that were, were classified the same. At the order to species rank, OTUs classified by both LSU and ITS were rare and differences between the markers were more common (Fig. 5F).

**Figure 5:**
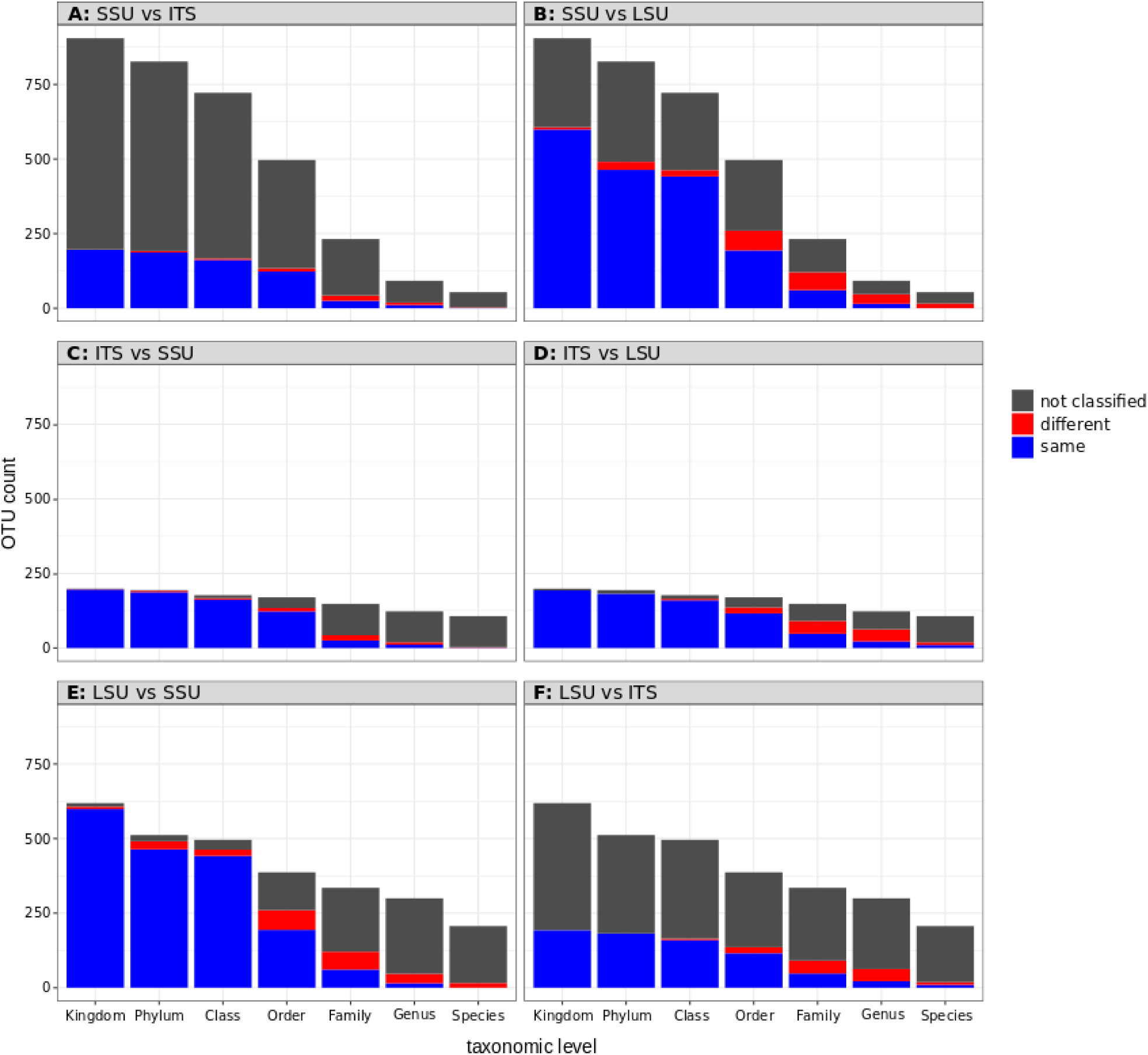
Agreement of classifications of all OTUs by the different regions. Each panel represents a comparison between two regions. Each set of stacked bars shows numbers of agreeing (blue), disagreeing (red) and unknown (gray) OTU classifications in the second region of the comparison compared to the first at each taxonomic level.

## DISCUSSION

Long sequencing reads have the potential to provide many benefits for DNA metabarcoding. These include taxonomic assignment of OTUs at lower taxonomic levels (Porter & Golding 2011, Franzén 2015), the use of homology-based classification and phylogenetic reconstruction (e.g., Tedersoo 2017), and higher sequencing quality for standard-length DNA barcodes in reference databases (Hebert 2017). Disadvantages of long reads include lower sequence quality (Glenn 2011, D’Amore and Ijaz 2016), a possible increase in the rate of chimera formation, and the fact that most bioinformatics tools are optimized for shorter reads. Here we produced DNA metabarcodes nearly twice as long as any used to date (ca. 4,500 bp), comprising the whole eukaryotic rRNA operon (SSU, ITS, LSU). We combined circular consensus sequencing with our newly developed bioinformatics pipeline and obtained error rates comparable to short-read Illumina sequencing (Glenn 2011, D’Amore and Ijaz 2016). The use of multiple markers allowed us to use the ITS region for OTU delineation (clustering) and automated species-level taxonomic classifications for environmental OTUs with both ITS and LSU sequences. Finally, the inclusion of the SSU rRNA gene into the analyses allowed us to classify OTUs that were not represented in ITS and LSU databases, including many fungi that belong to basal lineages and are common in freshwater habitats (Rojas-Jimenez 2017, Wurzbacher 2016).

### Challenges of long reads

A significant challenge in using longer reads for DNA-metabarcoding of mixed samples is the fact that most bioinformatics tools have been designed for the analysis of short sequences (typically 200-600 bp). Although we obtained very high-quality CCS reads, the higher indel rate and accumulation of errors in long reads requires analyses that differ from that of more commonly used sequencing platforms like Illumina. For example the clustering algorithm applied by swarm (Mahé 2015) relies on a low total number of errors per sequence (ideally 1 error). In long sequences, even with low error rates, the total number of errors are higher, makeing it unfeasible to use this algorithm. Other widely used clustering tools like uclust (Edgar 2012) or vsearch use heuristics to choose starting points for clustering. Reads are first de-replicated and those with the most identical copies are used as cluster starting points. This could not be applied to our data set because the comparably high nucleotide deletion rate and the long read length made almost all reads unique.

In the future it might be beneficial to develop specialized software for clustering and correcting PacBio long range amplicons. Here we used heuristic clustering starting with high quality reads and with a high similarity threshold (99%), and a consecutive consensus calling for correction of random sequencing errors. This also gave us clusters of highly similar sequences, that we could use for chimera detection and OTU clustering instead of the groups of identical reads resulting from de-replication, that are normally used for these steps.

One of the problems in any study applying PCR to mixed samples is chimera formation. Our comparison of *de novo* and reference-based chimera detection found them to produce the same classifications in > 98% of cases. This indicates that *de novo* chimera classification in our long-read pipeline provided a good estimate of chimera formation rate and is suitable for data sets where no complete reference database is available. We can therefore be confident in our results for the environmental samples, even where no reference sequences were available in databases. Interestingly, a manual inspection of coneicting read assignments in the independent clustering of the different regions (data not shown) found a few cases (9 of 6,585 reads in the one mock community sample) of chimeras that could not be detected. Neither reference-based nor *de novo* approaches detected these chimeras because 3’ and 5’ ends were both from the same species, and only the central section originated from a second species. Most chimera detection software, including vsearch, model chimeras from two origins i.e., different 3’ and 5’ ends, but not more. These methods would then fail to identify chimeras if the 3’ and 5’ ends are from the same species and a second species is in the middle, as we observed. Although this was very rare in our data (0.1% of reads investigated), it created small OTUs made up almost entirely of these complex chimeras in our mock community (OTU 14, 16 and 17, see Table 4). As a general rule, chimeras are most likely to be found associated with the most frequent sequences in a PCR sample (e.g. Sommer 2013) and this is also true for the complex chimeras we observed here. In fact, all three chimeric OTUs found in our mock community involved the species with the most read abundance, *Metschnikowia reukaufii*. DECIPHER (Wright 2012) is one tool that may detect these chimeras, but requires a complete reference database of possible parent sequences and is therefore unsuitable for use with environmental samples (for which reference sequences are difcult to obtain) and long reads.

We also attempted to minimize chimera formation in the laboratory, by exploring the ineuence of reduced PCR cycle numbers, emulsion PCR, and template concentration. Although we were initially concerned that our *ca.* 4,500 bp amplicon length would lead to higher chimera formation rates during PCR, the mock community sample that was amplified with the highest cycle number (30) formed chimeras at a rate within the range reported by short-read studies (*ca.* 4-36%; Qiu 2001, Ahn 2012). We observed reduced chimera formation with fewer cycles which is also consistent with short-read studies (Qiu 2001, Lahr and Katz 2009, D’Amore and Ijaz 2016). Unlike other studies (Lahr and Katz 2009, D’Amore and Ijaz 2016) we did not fnd a notable ineuence of DNA template concentration in our samples, possibly because at 18 cycles all reactions were still in the exponential phase, before depletion of reagents (see below). Chimera formation rates in our mock community may underestimate rates in environmental samples because the lower species richness in the mock community may have led to reduced chimera formation (Fonseca 2012). However, the chimera rate detected by *de novo* chimera detection in our environmental data was < 1%, i.e., even lower than the *de novo* detection rate in the less diverse mock community samples. Chimera formation occurs primarily during the saturation phase of a PCR, when a large amount of PCR product has accumulated and the template:primer ratio increases (Judo 1998). For a given cycle number, the amount of accumulated product may differ between the environmental and mock community samples, because although a similar amount of template DNA was used in mock community (8 ng) and environmental (10 ng) samples, the amount of template available for primer binding might be lower in the latter because they also contain non-fungal DNA. Environmental samples may also contain more PCR inhibitors (Schrader 2012), which would reduce PCR efciency and delay the saturation phase to a higher cycle number in environmental samples compared to the mock community. Optimization of DNA extraction and amplification could make lower PCR cycle numbers feasible and thus further reduce the problem of chimera formation. Our emPCR results also indicate that this might be a promising way of reducing chimera formation when more PCR cycles are required.

### Classification

Although the ITS region has been proposed as a standard barcode for fungi (Schoch 2012) other regions of the rRNA operon remain popular choices as fungal barcodes (Stielow 2015, Roy 2017, Wurzbacher 2016). Compared to rRNA genes, ITS1 and ITS2 often exhibit higher interspecific variability and thus can provide greater species delineation power (i.e., more OTUs) than SSU and (in most fungal groups) LSU (Schoch 2012). Indeed we found that isolate species of the same genus (Cladosporium) and even from the same order (Hypocreales) and sub-division (Pezizomycotina) could not be separated by the SSU (Fig. 3), and that the use of ITS resulted more often in classification to species level than SSU and LSU in Dikarya (Fig. 4). At the same time, the often higher variability of ITS also means that for new species that are not represented in the database it can be more difcult to fnd comparable sequences and thus to identify them to any level. In these cases, longer sequencing reads that include more conserved regions with a stable evolutionary rate are likely to be helpful in making classifications based on sequence similarity as we did here or, by phylogenetic methods (e.g., Tedersoo 2017). The phylum Chytridiomycota, which is often found in aquatic environments and was highly abundant in our environmental samples, is underrepresented in sequences databases (Frenken 2017). We observed many OTUs from this phylum that could not be classified with the ITS at all, while the SSU provided at least class or family rank classifications and the LSU often even provided classifications at species rank (Fig. 4).

For the classification of the mock community, the different degrees of taxonomic resolution provided by the different markers were clear. The mock community consisted of species that are represented in the reference databases with sequences that were identical or very similar to the sequence that we found. In these cases, ITS was a superior marker region, since its greater variability allowed for higher resolution classification. While almost all classifications were correct, those obtained for ITS went down to at least family rank in all cases, and even to species rank for a third of the OTUs. LSU and SSU both provided far fewer specific classifications. Using the LSU marker, species levels classifications could be obtained for some OTUs, but others were only classified to higher taxonomic ranks (up to kingdom). Using the SSU marker, classification results were obtained between the ranks of order and family. In our environmental samples, the disadvantage of ITS becomes clear. If no closely related reference sequence was available, sequence similarity to any sequence in the database was too low to classify the sequence even to a higher taxonomic rank. In these cases, SSU and LSU markers provided at least classification at family or class level, while many OTUs stayed completely unclassified with the ITS.

The independent clustering of the different regions (SSU, ITS1, 5.8S, ITS2 and LSU) of the rRNA operon (Fig. 2) also showed the higher resolution of ITS1 and ITS2, which were the only regions that separate almost all species from each other. On the other hand, for *Metschnikowia reukaufii* they formed multiple clusters for one species. This is most likely the result of high variability of rRNA operon copies in *Metschnikowia* (Sipiczki 2013, Lachance 2003) in combination with the short ITS1 and ITS2 sequences (70 bp and 75 bp, respectively) which mean that very few (3) differences already constitute an identity difference of 3%.

### Classification conficts and synergies

The coneicts we observed between classifications based on different marker regions and databases can provide insights into a number of interesting problems. In some cases, they may either represent uncertainty in classification using at least one of the markers, or genuine chimeric reads. In other cases they may highlight incompatibility between the taxonomies used by the databases, or even errors in the databases (see also Nilsson 2006). Many coneicts resulted from differences in naming convention and taxonomic placement in the different databases. Multiple OTUs were classified with LSU and the RDP database to the more recently defined orders Rhizophydiales (Letcher 2006) and Lobulomycetales (Simmons 2009), but were classified with SSU and the SILVA database as Chytridiales, the older classification for these new orders. A similar effect can be seen for the orders in the class Agaricomycetes. Three OTUs were assigned to the family Lachnocladiaceae which belongs to the order Russulales according to SILVA and to Polyporales according to RDP. Finally, one OTU was assigned to the genus *Jahnoporus* using the LSU marker. According to the RDP database this genus belongs to the order Russulales while in SILVA it belongs to the order Polyporales. Other coneicts showed that minor problems in the databases can lead to major differences in classification. In our environmental data, several high (read) abundance OTUs were classified as Chytridiomycota with SSU but as Blastocladiomycota with LSU. Closer inspection of the LSU alignments indicated that for many of these OTUs, only the second best hit was to a Blastocladiomycota, while the best match was, in fact, *Rhizophlyctis rosea*. The latter is a Chytridiomycota, but has no classification beyond kingdom in the RDP database file we used and was thus ignored for classification. In addition, the second best hit which was used for classification is to a sequence from the genus *Catenomyces* which belongs to the phylum Blastocladiomycota according to RDP, but according to SILVA belongs to the phylum of Chytridiomycota. Thus a minor error in the database file, in combination with inconsistencies in the taxonomy used by different databases, can lead to completely different classifications when using different markers.

These coneicts in classification clearly highlight problems with the databases, but classifications using three different markers from the same molecule, as obtained from the full rRNA operon, can help us to evaluate how confident we can be in our classification. A classification that is supported by three markers, with largely independent databases, can be considered more trustworthy than one that is only supported by one, or even shows coneicts when using different 685 markers. In addition, long DNA barcodes could be used to create synergies between the databases and to support short read studies. For example, if a sequence was classified to the same family by SSU (SILVA) and LSU (RDP), the ITS region could be added to the Unite database (even if it is not classified to the species level) to help future studies that use ITS markers. The possibility to sequence SSU, ITS and LSU at the same time therefore offers the opportunity to contribute to different databases in parallel, with the future potential to generate a new reference data set with nearly full-length rRNA operon sequences.

### Conclusions

We used a DNA metabarcode nearly twice the length of any used to date and created a long-read (*ca*. 4,500 bp) bioinformatics pipeline that results in rates of sequencing error and chimera detection that are comparable to typical short-read analyses. The approach enabled the use of three different rRNA gene reference databases, thereby providing significant improvements in taxonomic classification over any single marker. While ITS is likely to remain a short-metabarcode region of choice for some time, a clear limitation of ITS is that its high variability, in combination with the incompleteness of databases, often lead to classification failing. In these cases, the other rRNA markers are beneficial. In particular, classification based on SSU or LSU were superior in more basal fungal groups. The universal nature of the rRNA operon and our recovery of >100 non-fungal OTUs indicate that the method could also be suitable for more general studies of eukaryotic biodiversity.

## ACKNOWLEDGEMENTS

We thank Lars Ganzert, Katrin Premke, and Robert Taube (IGB) for help with field sampling, Keilor Rojas and Silke Van den Wyngaert (IGB) for providing isolates, Christian Wurzbacher (Univ. Gothenberg, now TU Munich) for providing primers, and Nicole Heyer and Simone Severitt (DSMZ) for help with sequencing. Research was partially funded by the Leibniz Association Pakt/SAW project “MycoLink” (SAW-2014-IGB-1).

