## Supplementary Materials for "Long-read DNA metabarcoding of ribosomal rRNA in the analysis of fungi from aquatic environments"

### Supplemental tables:

Supplemental Table 1. 16-mer barcodes
Supplemental Table 2. PCR details (template, cycle numbers), emPCR.
Supplemental Table 3. Library and barcode setup
Supplemental Table 4. OTU table

### Supplemental info 1: DNA extraction of isolates

For the isolates extracted with the Qiagen Dneasy Plant Mini Kit (Qiagen, Hilden, Germany), cultures were maintained on 2 % malt extract agar (MEA, Carl Roth, Karlsruhe, Germany) and mycelia harvested directly from MEA plates. Fresh mycelium was either ground in lysis buffer AP1, using 1.5-ml micropestles (Sigma-Aldrich, Darmstadt, Germany), or was first lyophilized, and 20 mg homogenized to a fine powder in 2-ml tubes containing ¼ tube volume of beads (1x 3 mm steel ball, 2x 2 mm glass beads, 1 mm Zirconium beads, Biospec, Bartlesville, USA) at 30 Hz for 2 min, using a Retsch mill (Retsch GmbH, Haan, Germany).. Next, 400 µl buffer AP1  and 4 µl  RNase A solution was added. For both methods, lysis was carried out for 1 hr at 65 °C. Extraction thereafter followed manufacturer's instructions.

For isolates extracted with the MasterPure™ Yeast DNA Purification Kit (Epicentre, Madison, US), cultures were maintained on 2 % MEA and mycelia were harvested directly from MEA plates. Genomic DNA was extracted in conjunction with the FastPrep 24 instrument (MP Bio, Eschweg, Germany) according to the manufacturer's instructions.

For the peqGOLD Tissue DNA Mini Kit (VWR, Vienna, Austria), isolates were maintained and genomic DNA extracted as described in Rojas-Jimenez et al. 2017. DNA was extracted from approx. 200 mg of fungal material following the manufacturer's instructions.

DNA was checked on a gel, and quantified in duplicate using a Qubit HS dsDNA Assay using the Qubit 2.0 system (Invitrogen, Carlsbad, USA). A mock community was formed from pooling DNA according to Table 1 (main document).

### Supplemental Info 2: emPCR

Emulsion PCR was performed over two amplification steps, using the Micellula DNA Emulsion Kit (Roblokon, Berlin, Germany), following manufacturer's instructions. Two steps were used to ensure sufficient PCR product for later sequencing as a single round of PCR yields a low level of product.

We used 20 ng of mock community DNA in a first (pre-PCR) 50 µl reaction. This amount was determined to give a sufficiently high input to populate enough water droplets in the emulsion with DNA template, while limiting their overpopulation (which would remove the advantage of an emulsion PCR).

Water-in-oil emulsions were prepared as follows: the PCR mixture was set up as a 50 µl reaction, following the standard Herculase II Fusion PCR reactions described in the main text, with the exception of a reduced BSA input of 0.025 µl per reaction. The emulsion was created by adding the aqueous reaction to 300 µl pre-cooled Oil Surfactant Mixture and vortexing the solution for 5 minutes at 4 °C, before being split equally (112 µl) across three PCR tubes.

The PCR conditions followed those given in the main text, for 25 cycles. Following the first pre-PCR, the split sample was re-pooled and the emulsion broken by the edition of 1 ml 2-butanol (Sigma-Aldrich), followed by a column clean-up (Micellula DNA Emulsion Kit) according to Manufacturer's instructions. The product concentration was checked with a Qubit HS dsDNA Assay. We performed a (scale-up) PCR using 2 ng of the first PCR product as template in a second emulsion PCR reaction to yield enough PCR product for sequencing. Primers were barcoded as described in supplementary Table 1, to allow for multiplexing before library preparation.

### Supplemental Info 3: Pipeline steps

#### getFullCls

For each pre-cluster, representative sequences get a classification. This can be either  i) CHIMERA if the reference base chimera detection called this as chimeric (Y) or possibly chimeric (?),  ii) UNKNOWN if the sequence was not called as chimeric, but not match to an isolate consensus sequence was found or iii) the name of species, this is given by the highest scoring match to a isolate consensus sequence for non-chimeric sequences.

#### fullMapping

Run blasr to map representatives of non-chimeric pre-clusters against the isolate consensus sequences. The following parameters are used: `-m 5` to get tabular output, `--bestn 50` to get a maximum of 50 hits for each query and `--minPctSimilarity 90` to get only hits with at least 90% identity.

**removeChimeraRef**

Run vsearch to remove chimeras with reference based approach. The `--uchime_ref` parameter is used to run the reference based chimera detection algorithm and the `--db` parameter to give the isolate consensus sequences selected by the getFullRef rule as reference sequenes.

**getFullRef**

For each isolate sample get the consensus sequence of the biggest pre-cluster. Will give an error if there are more than one pre-cluster with 10 or more reads for one sample. Compares sequences for replicates of each species (if available) and writes a warning to the log file if a difference is encountered.

**getCorrectCls**

Get “correct” classifications for reads in each OTU according to mappings to isolate consensus sequences. Each OTU might have multiple species with read numbers listed.

**classifyLSU**

Classify OTUs by matches of LSU sequences to the RDP LSU database. See main methods section for details and parameters.

**alignToRdp**

Run lambda to get local alignments of each OTU LSU representative to the RDP LSU database. Lambda is run with the following parameters: `--output-columns "std qlen slen"` to get query length and subject length along with the default columns in the output table (these are used for coverage computation later), `-p blastn` to run in blastn (nucleotide vs nucleotide) mode, `-nm 5000` to get more matches per query sequence (this is important due to the high number of similar sequences in the database), `-b -2` to the square root of the query length as width for the banded alignment optimization (because higher indel rate and, even more important, uneven insertion/deletion ratio cause the alignment to leave the band), `-x 40` to reduce the x-drop value which can cause alignments to be terminated prematurely and `-as F` to disable adaptive seeding which normally is used to reduce number of hits. Parameters were optimized to allow for alignments for all mock community species.

**classifySSU**

Classify OTUs by matches of SSU sequences to the SILVA database. See main methods section for details and parameters.

**alignToSilva**

Run lambda to get local alignments of each OTU SSU representative to the SILVA database. Lambda is run with the following parameters: `--output-columns "std qlen slen"` to get query length and subject length along with the default columns in the output table (these are used for coverage computation later), `-p blastn` to run in blastn (nucleotide vs nucleotide) mode, `-nm 20000` to get more matches per query sequence (this is important due to the high number of similar sequences in the database), `-b -2` to th square root of the query length as width for the banded alignment optimization (because higher indel rate and, even more important, uneven insertion/deletion ratio cause the alignment to leave the band), `-x 30` to reduce the x-drop value which can cause alignments to be terminated prematurely and `-as F` to disable adaptive seeding which normally is used to reduce number of hits. Parameters were optimized to allow for alignments of all mock community species.

**classifyITS**

Classify OTUs by matches to the UNITE database. See main methods section for details and parameters.

**alignToUnite**

Run lambda to get local alignments of each OTU representative to the UNITE database. Lambda is run with default parameters except for: `--output-columns "std qlen slen"` to get query length and subject length along with the default columns in the output table (these are used for coverage computation later) and `-p blastn` to run in blastn (nucleotide vs nucleotide) mode.

**otuCluster**

Run vsearch to cluster OTUs at 97% identity threshold. The following parameters are used for vsearch: `--cluster_size` to choose cluster seeds by descending pre-cluster size (according to size annotation), `--relabel otu` to name OTUs with out and running number instead of using the first sequence as a name, `--sizein --sizeout` to read and write size annotation, `--iddef 0` to use the identity definition of CD-Hit (see vsearch manual), `--id 0.97` to use 97% identity threshold and `--minsl 0.9` to only accept alignments of at least 90% coverage for similarity computation. In addition `--centroids` is used to output a representative centroid sequence for each OTU.

#### itsx

Run ITSx to separate the different regions of the rRNA operon. The following parameters were set for ITSx: `-t .` to use HMM models from all available taxonomic groups, `--save_regions SSU,ITS1,5.8S,ITS2,LSU` to save separate files for all different regions of the rRNA operon, `--complement F` to not allow reverse-complement detection (all sequences were orientated in forward direction in the primerFilter rule), `--partial 500` to allow for partial matches (we do not have complete SSU and LSU sequences in the amplicon) and `-E 1e-4` to allow for HMM hits with slightly lower e-values (this was optimized to make sure that all rRNA operons in the mock community species were recognized).

#### removeChimera

Run vsearch to remove chimeras in *de novo* mode. Vsearch is run with the `-uchime_denovo` parameter. All parameters for the chimera detection algorithm are left at default values. Vsearch automatically uses size annotations form the pre-clustering step for its greedy algorithm.

#### preCluster

Run vsearch to create pre-clusters at 99% identity threshold. The following parameters are used for vsearch: `--usersort --cluster_smallmem` to choose cluster seeds in the order the sequences are sorted in the input file, `--relabel` to name pre-clusters according to the given scheme instead of using the first sequence as a name, `--sizeout` to add size annotation to the output, `--iddef 0` to use the identity definition of CD-Hit (see vsearch manual), `--id 0.99` to use 99% identity threshold and `--minsl 0.9` to only accept alignments of at least 90% coverage for similarity computation. In addition `--consout` is used to generate consensus sequences for each pre-cluster.

#### prepPrecluster

Reads are sorted by descending mean quality (see qualityFilter rule for computation). This helps to use high quality reads as cluster seeds for pre-clusters in the next step.

#### filterPrimer

Primers are found and cut with cutadapt. Cutadapt is run with default parameters except for `--trimmed-only` to only retain reads were the primer was found and `-O 10`. Cutadapt is configured to search for both the forward and the reverse primer at the start of the sequence. For sequences where the forward primer was found, the reverse-complemented reverse primer is search at the end of the sequence with an additional run of cutadapt. Accordingly for sequences where the reverse primer was found, the reverse-complemented forward primer is searched at the end of the sequence with another run of cutadapt. In the end sequences with forward-reverse primer combination and reverse-complemented sequences with reverse-forward primer combination are concatenated into one file.

**windowQualFilter**

For overlapping windows of size 8 the mean error rate is computed from the Phred scores with the same formula as in the qualityFilter rule (except that S is the substring in the windiw instead of the whole sequence). If any window in a sequence has a mean error rate of 0.9 or higher the sequence is removed.

**qualityFilter**

Mean error rate per sequence is computed from the Phred score given in the fastq file with the formula: $\frac{\sum 10^{-q/10}}{length(S)}$ with q being the quality values of sequence S. Sequences with an error rate of 0.4% or more are written to a separate file (not further used).

**lengthFilter**

Sequences with length above 6,500 or below 3,000 are printed to separate files (not used further).

**filterSilva**

Filter SILVA sequences by the quality and pintail (chimera probability) values given in the database. Only sequences with a quality value of at least 85 and pintail value of at least 50 are retained.
